# A versatile dual reporter to identify ribosome pausing motifs alleviated by translation elongation factor P

**DOI:** 10.1101/2024.08.03.606492

**Authors:** Urte Tomasiunaite, Tess Brewer, Korinna Burdack, Sophie Brameyer, Kirsten Jung

## Abstract

Protein synthesis is influenced by the chemical and structural properties of the amino acids incorporated into the polypeptide chain. Motifs with consecutive prolines can slow down translation speed and cause ribosome stalling. Translation elongation factor P (EF-P) facilitates peptide bond formation in these motifs, thereby alleviating stalled ribosomes and restoring regular translational speed. Ribosome pausing at various polyproline motifs has been intensively studied using a range of sophisticated techniques, including ribosome profiling, proteomics, and *in vivo* screenings with reporters incorporated into the chromosome. However, the full spectrum of motifs which cause translational pausing in *Escherichia coli* has not yet been identified. Here we describe a plasmid-based dual reporter for rapid assessment of pausing motifs. This reporter contains two coupled genes encoding mScarlet-I and chloramphenicol acetyltransferase to screen motif libraries based on both bacterial fluorescence and survival. In combination with a diprolyl motif library, we use this reporter to reveal motifs of different pausing strengths in an *E. coli* strain lacking *efp.* Subsequently, we use the reporter for a high-throughput screen of four motif libraries, with and without prolines at different positions, sorted by fluorescence-associated cell sorting (FACS) and identify new motifs that influence translational efficiency of the fluorophore. Our study provides an *in vivo* platform for rapid screening of amino acid motifs that affect translational efficiencies.

## Introduction

The regulation of protein synthesis is critical for the maintenance of physiological balance and the survival of organisms. A number of factors can modulate translational efficiency, including charge-specific interactions between the ribosome peptide exit tunnel and the nascent chain^1–3^, mRNA secondary structures^4–7^, tRNA abundance and modification^8–10^, and codon selection^11,12^. It has been widely demonstrated that certain combinations of amino acids, in particular stretches of polyprolines, have the ability to negatively impact translational efficiency and in certain instances cause the ribosome to stall^13–16^. The profound negative effect of proline on translation rate can be attributed to its unique structural nature, which includes a pyrrolidine ring that renders it conformationally rigid. This ultimately results in proline being a poor donor and poor acceptor during the transpeptidation reaction^17–20^. Despite the fact that proline is a challenging factor during translation, diprolyl motifs (XPPX) are very common in bacterial proteomes. In *Escherichia coli*, there are 2,184 of these motifs (0.51 motif/protein), in *Salmonella enterica* 2,324 motifs (0.51 motif/protein), in *Pseudomonas aeruginosa* 4,074 motifs (0.73 motif/protein) and *Streptomyces coelicolor* even contains 8,719 motifs (1.08 motif/protein)^21,22^. The prevalence of these motifs is not limited to bacterial proteomes — they are also common among the proteomes of eukaryotes^23^. The ubiquity of these motifs in the proteomes of organisms underlines their functional and regulatory importance to shape protein structure^24,25^, adjust copy number^21^ and incorporate transmembrane proteins^26^.

The biosynthesis of proteins containing polyproline motifs is mediated by distinct translation factors that have evolved in organisms from all domains of life. While eukaryotes and archaea encode initiation factor 5A (e/aIF5A)^27^, bacteria have evolved elongation factor P (EF-P), which reduces ribosomal stalling by assisting during the translation of polyproline motifs^14,15^. The EF-Ps of numerous bacterial species are dependent on unique post-translational modifications (PTM), which are necessary to improve their activity^28–35^. In contrast, certain EF-Ps, particularly from the subfamily with the activation loop PGKGP (Pro-Gly-Lys-Gly-Pro), exhibit full functionality in the absence of PTMs, although with reduced levels of activity in *E. coli*^22,36^. Intriguingly, our recent findings even demonstrate that some unmodified EF-Ps, which are barely active in *E. coli*, can be engineered functional in *E. coli*^36^. This suggests that full EF-P functionality is not exclusively dependent on PTM, as previously assumed. Notably, we observed variations in the efficacy of distinct polyproline motif rescue among modified and unmodified EF-Ps, suggesting that the underlying general mechanisms of protein synthesis are not yet fully understood and require further investigation.

A number of studies have employed a range of techniques, including ribosome profiling, mass spectrometry, in *vitro* and in *vivo* reporter systems, to identify motifs associated with different degrees of ribosome pausing^14,16,37–41^. The majority of these studies focuses on motifs which contain at least two prolines, as EF-P is well known to support their synthesis during the transpeptidation reaction^16,38^. The literature on this topic is limited with regards to motifs that contain a single proline or even no proline, which rescue is dependent on a functional EF-P^37,42^. The existing knowledge gaps highlight the need for simple, accessible tools that can be utilized by every laboratory at a low cost, with a minimal input of materials to investigate translational dynamics.

In this study, we present a user-friendly, plasmid-based reporter system for investigating translational efficiencies of amino acid motifs. Translationally coupling^43^ the genes encoding the fluorophore mScarlet-I and chloramphenicol acetyltransferase, enhances screening capacity, allowing dual output measurements of cell fluorescence and viability. High-throughput screening demonstrates that this system effectively identifies unusual sequence motifs, particularly those lacking proline, which cause reduced translation rates.

## Results

### A plasmid-based in *vivo* dual reporter system enables ribosome pausing measurements in a Δ*efp* mutant

To facilitate rapid evaluation of which amino acid motifs can influence translational efficiency, we established a plasmid-based reporter system. Motifs which are known to cause various levels of ribosome pausing were chosen for initial experiments: RPAP (Arg-Pro-Ala-Pro) – represented no pausing strength, TPPP (Thr-Pro-Pro-Pro) – represented middle pausing strength, and RPPP (Arg-Pro-Pro-Pro) – represented strong pausing strength^14,38,41^ (**Figure 1A**). To identify differences in translational efficiency resulting from distinct sequence motifs, an *E. coli* BW25113 mutant lacking *elongation factor P* (Δ*efp*) was transformed with these reporter plasmids. The initial reporter design consisted of an antibiotic resistance cassette (*chloramphenicol acetyltransferase, cat*) with motifs integrated at the N-terminus (**Supplementary Figure 1A,** left panel). This approach allowed estimation of the translational efficiencies of *cat* influenced by the motif sequence, based on bacterial survival in the presence of chloramphenicol. A 24-hour incubation in media containing serial dilutions of chloramphenicol demonstrated differences in bacterial survival only between clones carrying the reporters RPAP_*cat* and RPPP_*cat,* particularly at chloramphenicol concentrations 3.4 μg/mL, 3.8 μg/mL and 4.3 μg/mL (**Supplementary Figure 1B-C**). However, no significant differences between the clones carrying the reporters RPAP_*cat* and TPPP_*cat* could be detected at any of the chloramphenicol concentrations tested (**Supplementary Figure 1C**).

**Figure 1.**
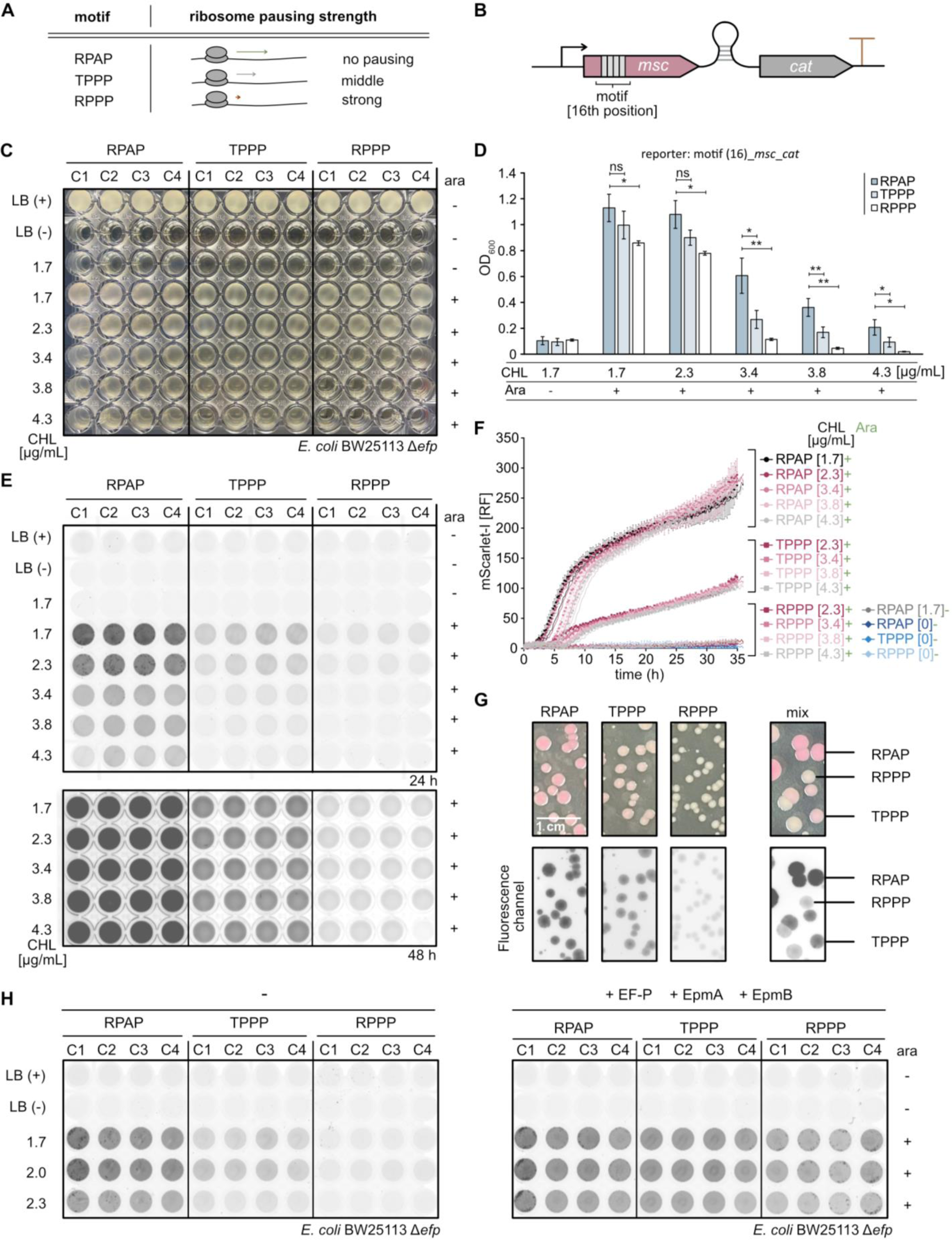
Development of a dual reporter system to screen for ribosome pausing motifs. **A** Schematic of known ribosome pausing strengths at motifs RPAP (Arg-Pro-Ala-Pro), TPPP (Thr-Pro-Pro-Pro) and RPPP (Arg-Pro-Pro-Pro) alleviated by elongation factor P^14,38,41^. **B** Construction of dual reporter for translational efficiency measurements. The reporter consists of a gene coding for fluorophore (mScarlet-I) and a resistance gene cassette (*chloramphenicol acetyltransferase, cat*), both linked by a weak translational coupling device^43^. **C-D** 96-well assay plate of the growth measurements (**C**) and growth quantification (**D**) of *E. coli* BW25113 Δ*efp*, transformed with the dual reporter plasmid pBAD24_motif_*mscarlet-I*_*cat*. **E** Fluorescence measurements of the 96-well assay plate 24 hours (upper panel) or 48 hours (bottom panel) after bacteria inoculation. *E. coli* BW25113 Δ*efp* was transformed with the dual reporter plasmid pBAD24_motif_*mscarlet-I*_*cat*. **F** Relative fluorescence (RF) measurements over time of *E. coli* BW25113 Δ*efp* with dual reporter plasmid pBAD24_motif_*mscarlet-I*_*cat*. The M9 medium was supplemented with different concentrations of chloramphenicol (CHL). mScarlet-I fluorescence signal was normalized to the measured optical density (OD_600_) and shown as RF. **G** Colony morphology and fluorescence analysis of *E. coli* BW25113 Δ*efp* with dual reporter plasmid pBAD24_motif_*mscarlet-I*_*cat*. Bacteria were plated on LB agar assay plates supplemented with 0.2 % (w/v) arabinose (Ara) and 3.4 μg/mL CHL. Nucleotides encoding the RPAP, TPPP, and RPPP motifs were incorporated into the fluorophore gene sequence starting from the sixteenth nucleotide position (16th). **H** EF-P complementation assay. Fluorescence measurements of the 96-well assay plate 30 hours after bacteria inoculation. *E. coli* BW25113 Δ*efp* was transformed with the dual reporter plasmid pBAD24_motif_*mscarlet-I*_*cat* (left panel) or with the dual reporter plasmid pBAD24_motif_*mscarlet-I*_*cat* and pBAD_*E. coli efp*_*epmA*_*epmB* (right panel). Error bars indicate the standard deviation (SD) of three independent biological replicates. Statistics: student’s unpaired two-sided t test (\*\*\*\**p* < 0,0001; \*\*\**p* < 0.001; \*\**p* < 0.01; \**p* < 0.05; ns *p* > 0.05). 1.7 μg/mL CHL and 0.2 % (w/v) Ara (RPAP vs TPPP: ns *p* = 0.178 [D]; RPAP vs RPPP: \**p* = 0.016 [D]); 2.3 μg/mL CHL and 0.2 % (w/v) Ara (RPAP vs TPPP: ns *p* = 0.055 [D]; RPAP vs RPPP: \**p* = 0.013 [D]); 3.4 μg/mL CHL and 0.2 % (w/v) Ara (RPAP vs TPPP: \**p* = 0.015 [D]; RPAP vs RPPP: \*\**p* = 0.008 [D]); 3.8 μg/mL CHL and 0.2 % (w/v) Ara (RPAP vs TPPP: \*\**p* = 0.009 [D]; RPAP vs RPPP: \*\**p* = 0.003 [D]); 4.3 μg/mL CHL and 0.2 % (w/v) Ara (RPAP vs TPPP: \**p* = 0.038 [D]; RPAP vs RPPP: \**p* = 0.012 [D]). C1-C4 – clone number.

We hypothesized that reducing *cat* expression would enhance screening sensitivity, making the translational efficiencies of the RPAP and TPPP motifs more distinguishable. As previously described^43,44^, a weak translational coupling of a fluorophore gene upstream of a resistance gene cassette can support attenuation of antibiotic resistance and enable fine-tuning of screenings. Consequently, we constructed a dual reporter plasmid encoding for super folded green fluorescent protein (sfGFP) and CAT with a weak translational coupling device^43^ connecting the genes (**Supplementary Figure 1A**, middle and right panel). Motifs were incorporated into the gene encoding the fluorophore (**Supplementary Figure 1A**, middle and right panel). Endpoint optical density quantification revealed that chloramphenicol influenced the survival of all clones tested with the dual reporter motif_*sfgfp_cat*, regardless of the specific motif present (**Supplementary Figure 1D**). Significant differences between all clones in media supplemented with 3.4 μg/mL or 3.8 μg/mL chloramphenicol were only detectable when motifs were at the sixteenth nucleotide position of the fluorophore sequence (**Supplementary Figure 1E**). This observation confirms the essentiality of the first fifteen nucleotides which are critical for the initial synthesis of the nascent peptide chain^45^.

To increase visibility on agar plates, we replaced *sfgfp* with *mscarlet-I*, since the fluorophore mScarlet-I changes the colony color on agar plates to bright pink as shown in earlier studies^46^ (**Figure 1B**). Endpoint quantification of optical densities demonstrated that all Δ*efp* cells containing reporters encoding RPAP, TPPP and RPPP motifs exhibited differences in survival at 3.4 μg/mL, 3.8 μg/mL and 4.3 μg/mL chloramphenicol concentrations (**Figure 1C and D**). In particular, cells carrying the reporter RPPP_*mscarlett-I*_*cat* struggled to survive at chloramphenicol concentrations of 3.4 μg/mL or higher (**Figure 1C and D**). These findings imply that the translational efficiency of different motifs can be measured with the dual reporter system based on bacterial survival. Fluorescence intensity measurements after 24 hours (**Figure 1E**, upper panel) were in line with the observations from the bacterial survival assay (**Figure 1C-D**). We also extended the incubation period to 48-hours to ensure an equal optical density of bacterial culture in all wells, thereby demonstrating that the low fluorescence intensities observed with the RPPP_*mscarlett-I*_*cat* (**Figure 1E**, upper panel) reporter were not caused by differential cell abundance (**Figure 1E**, bottom panel).

We monitored relative fluorescence intensities for 35 hours in M9 minimal medium to investigate translational efficiencies over time (**Figure 1F**). The Δ*efp* strain with the reporter RPAP_*mscarlett-I*_*cat* exhibited the greatest relative fluorescence (RF) intensities, reaching a range of 200-300 RF between 25 and 35 hours (**Figure 1F**). The initial ten hours of monitoring revealed the impact of the supplemented chloramphenicol concentrations on the relative fluorescence (**Figure 1F**). The Δ*efp* strain containing the reporter TPPP_*mscarlett-I*_*cat* exhibited intermediate fluorescence intensities, reaching a range of 80-110 RFU between 25 and 35 hours (**Figure 1F**). Here, the impact of the supplemented chloramphenicol concentrations on the relative fluorescence curves was detected at the time interval of four to fourteen hours (**Figure 1F**). The low relative fluorescence intensities observed with strains containing the reporter plasmid RPPP_*mscarlett-I*_*cat,* were consistent with the endpoint fluorescence intensity measurements in **Figure 1E**. Regardless of the motif present, fluorescence was consistently low in the absence of arabinose and chloramphenicol, indicating low levels of leaky expression (**Figure 1F**).

To assess the reporter’s suitability for plate-based screens, we examined the growth of the Δ*efp* strain carrying reporters with RPAP, TPPP, or RPPP motifs on LB agar plates (**Figure 1G**). Preliminary analysis showed that LB agar plates with 3.4 μg/ mL chloramphenicol exhibited the most significant differences in colony morphology and fluorescence between cells containing reporters with either the RPAP or RPPP motifs **(Supplementary Figure 2A-C**).

After a 120-hour incubation period at room temperature, the colonies of Δ*efp* with RPAP_*mscarlett-I*_*cat* exhibited pink color and the largest colony diameter (**Figure 1G**). In contrast, Δ*efp* colonies expressing RPPP_*mscarlett-I*_*cat* exhibited yellow-white color and smaller colony diameters. In line with our measurements, Δ*efp* with TPPP_*mscarlett-I*_*cat* exhibited a light pink color and an intermediate colony diameter (**Figure 1G**). When EF-P and its modification machinery were overproduced in the Δ*efp* strain, translation of the fluorophore was rescued for all motifs (**Figure 1H**). This finding suggests that the observed reduction in fluorescence in the RPPP and TPPP reporters in Δ*efp* can be attributed to ribosome pausing events caused by the motifs in the absence of EF-P.

Taken together, our results demonstrate that the strength of ribosome pausing at amino acid motifs, can be effectively measured using a dual reporter system. Overall, these data show that mScarlet-I is an appropriate choice for the dual reporter system, enabling sensitive colony color contrast- and fluorescence-based screenings on plates to assess ribosome pausing strengths at specific motifs.

### Screening of libraries with diprolyl motifs demonstrates variation in translational efficiencies

We used plate-based screening of a diprolyl library (XPPX, X-Pro-Pro-X) in Δ*efp* to assess the efficacy of our dual reporter for large-scale translational efficiency measurements. We used a three-step protocol to generate the XPPX library, as illustrated in detail in **Figure 2A**. First, the dual reporter plasmid was amplified using primers with complementary overhangs. The forward primer was designed to be degenerate, incorporating all possible nucleotide (N) combinations between the two CCG codons coding for prolines (NNN-CCG-CCG-NNN) in the motif (**Figure 2A**). This strategy enables the incorporation of any amino acid in the positions upstream and downstream of the two prolines in the motif XPPX. Second, the *E. coli* MC1061 strain was transformed with the linearized XPPX library plasmids, allowing the library to circularize^43,44,47^. Third, the circularized XPPX library was isolated and the Δ*efp* mutant was transformed with the library (**Figure 2A**). The bacterial library populations exhibited diverse colony morphologies after plating on assay plates (**Figure 2B**). Qualitatively, we found that the colonies with the biggest diameter had a deep pink color, while the smaller colonies tended to be lighter pink or white-yellow (**Figure 2B**). The fluorescence analysis confirmed our previous observation that increasing levels of pink pigment resulted in higher fluorescence (**Figure 1G** and **Figure 2B**). These observations confirm that the amino acids which surround the two prolines in a diprolyl motif determine its translational efficiency, in line with published studies^16,21,39^.

**Figure 2.**
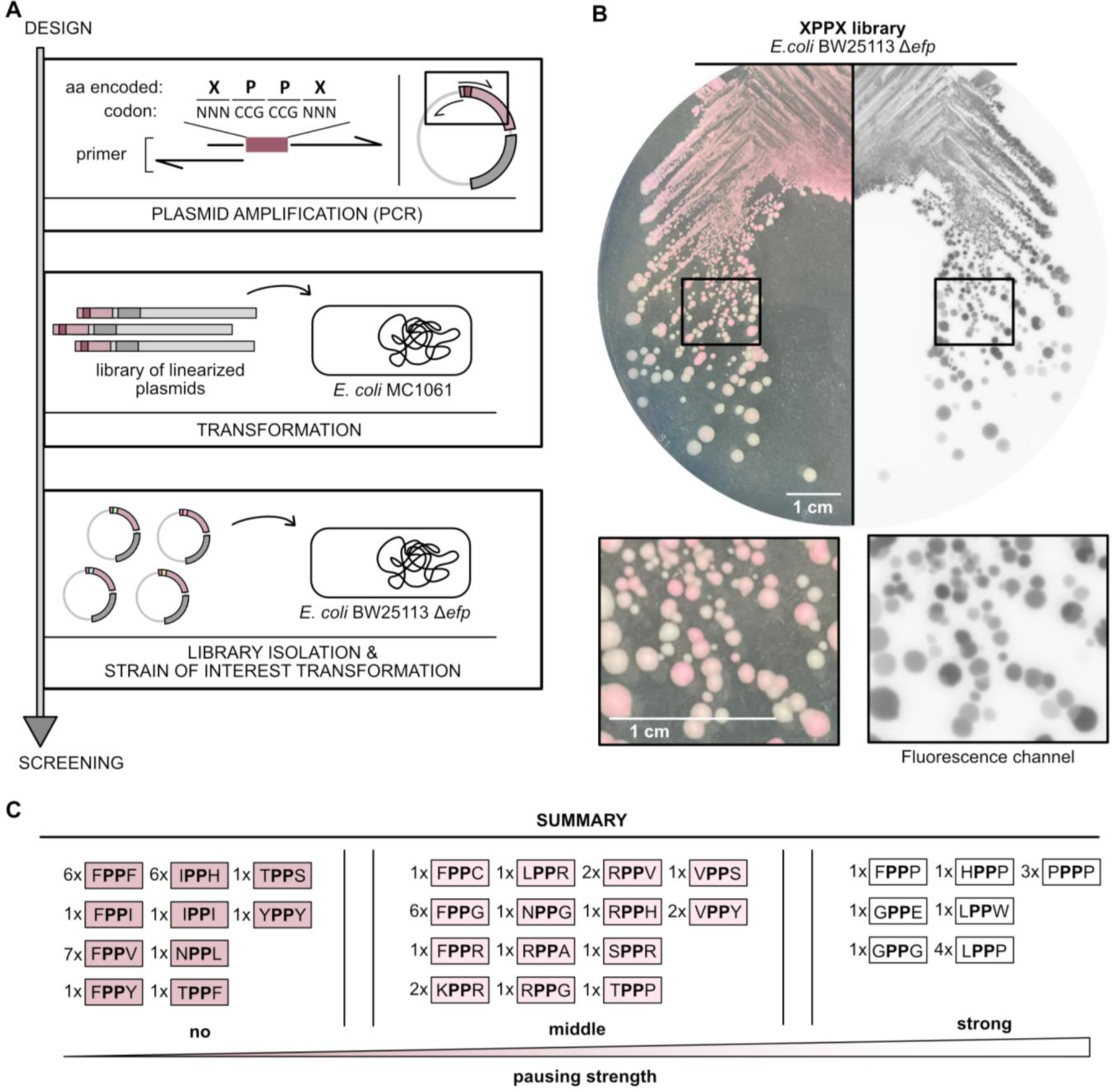
XPPX library screening on agar assay plates distinguishes motifs with different pausing strengths. **A** Graphical workflow to generate the XPPX (X-Pro-Pro-X) library (X represents all possible amino acids, N – all possible nucleotides). **B** Colony morphology and fluorescence analysis of *E. coli* BW25113 Δ*efp*, transformed with dual reporter plasmid pBAD24_XPPX_*mscarlet-I*_*cat* encoding the XPPX library. The images in the rectangles are shown enlarged under the agar plate. **C** Summary of the sequencing result of 60 clones that were picked according to their fluorescence intensity and viability. Motifs identified in pink colonies with high fluorescence intensities and large colony diameters were classified as those causing no pausing strength. Motifs observed in light pink colonies with intermediate fluorescence intensities and intermediate colony diameter were categorised as those causing middle pausing strength. Motifs identified in white-yellow colonies with low fluorescence intensities and small colony diameters were grouped as those causing strong pausing strength. The numbers indicate the count of plasmids found, that contain the indicated motif. aa – amino acid.

As a first pass, we manually screened sixty clones by grouping them into three categories based on colony color (pink, light pink and white-yellow colonies) and determined their motifs by Sanger sequencing (**Figure 2C**). We found some motifs overlapped in adjacent categories (for example between pink and light pink colonies) but found no overlaps between the pink and white-yellow color categories. Overlapping motifs were re-categorized into the category associated with weaker effects on translational efficiency. Altogether, we identified twenty-six sequences which had a marginal impact on the translational efficiency of the fluorophore (colonies with pink color), twenty-two sequences which caused moderate decreases in translational efficiency (colonies with light pink color), and twelve sequences which caused a strong decrease in translational efficiency (colonies with white-yellow color) (**Figure 2C**). Interestingly, motifs encoding FPPX were found across all categories of translational efficiency (X indicates any amino acid). When encoded in the FPPX motif, phenylalanine, isoleucine, valine and tyrosine (FPPF, FPPI, FPPV and FPPY) had a marginal influence on translational efficiency of the fluorophore. Cysteine, glycine and arginine (FPPC, FPPG and FPPR) caused a moderate decrease in translational efficiency, and proline (FPPP) had the strongest effect. These results indicate that the amino acid downstream of the two prolines in a diprolyl motif critically determines translational efficiency, consistent with previous findings^21^.

Together, these results demonstrate that our dual reporter can differentiate the pausing strength of diprolyl motifs in a Δ*efp* strain. We show that the strength of ribosome pausing can be assessed on agar plates supplemented with chloramphenicol by analyzing viability and the fluorescence of

### High throughput screening uncovers new motifs that reduce the translational efficiencies

To assess the reporter’s capability for high throughput translational efficiency measurements, we expanded our investigations using flow cytometry. To enhance the diversity of motifs we could analyze, we designed additional libraries with either a fixed CCG codon coding for a single proline (XPXX, NNN-CCG-NNN-NNN; XXPX, NNN-NNN-CCG-NNN) or completely randomized codons (XXXX, NNN-NNN-NNN-NNN). We sorted cells containing these plasmid libraries (about one million cells per library) with fluorescence-associated cell sorting (FACS) and analyzed the sorted cells with Illumina high throughput sequencing. In a pre-analysis, samples were treated with high temperatures (fifteen minutes at 95 °C, **Figure 3A**) to distinguish viable cells (P2) from non-viable cells (P1) with flow cytometry. As a first pass at identifying sequences which influence translational efficiencies, we calculated the log fold enrichment of particular motifs in the no/low fluorescence FACS bin versus the high fluorescence bin across multiple independent libraries (**Figure 3B**). We found that while generally the same motifs had similar values across different libraries, there was some variation (e.g. while RPPG was depleted in the no/low fluorescence bin in the XXXX, XPXX, and XXPX libraries, it was slightly enriched in the XPPX library, **Figure 3B**). Furthermore, we also detected variation in mScarlet-I expression by microscopy among cells carrying the same reporter plasmid (**Supplementary Figure 3A-C**). Thus, the appearance of motifs in both FACS bins could reflect heterogeneity in plasmid-based fluorophore expression, a phenomenon also reporter in earlier studies ^48,49^ (**Supplementary Figure 3D**). Ultimately, analyzing the relationship between both bins provides the most precise measurement of translational efficiency. To validate our results, we compared the top twenty motifs with the lowest and the highest translational efficiencies with predicted ribosome pausing strengths associated with distinct motifs from earlier studies^21^ (**Figure 3B**). We found good agreement between predicted pausing strengths and our results, indicating that our system is generally suitable for assessing translational efficiencies, despite variability in plasmid copy numbers within the bacterial population (**Supplementary Figure 3D**). Furthermore, while our analysis primarily focused translational efficiency at the amino acid level, it is likely that regulatory mechanisms at the codon level also play a role, as described in previous literature^41^. For example, several unique codon combinations were present for the amino acid motif RPPP in one of our libraries (**Supplementary Figure 4**). Differences in translational efficiency of different codon combinations which code for the same amino acid motif could also cause broad distributions of certain motifs across two distinct bins (**Figure 3B**).

**Figure 3.**
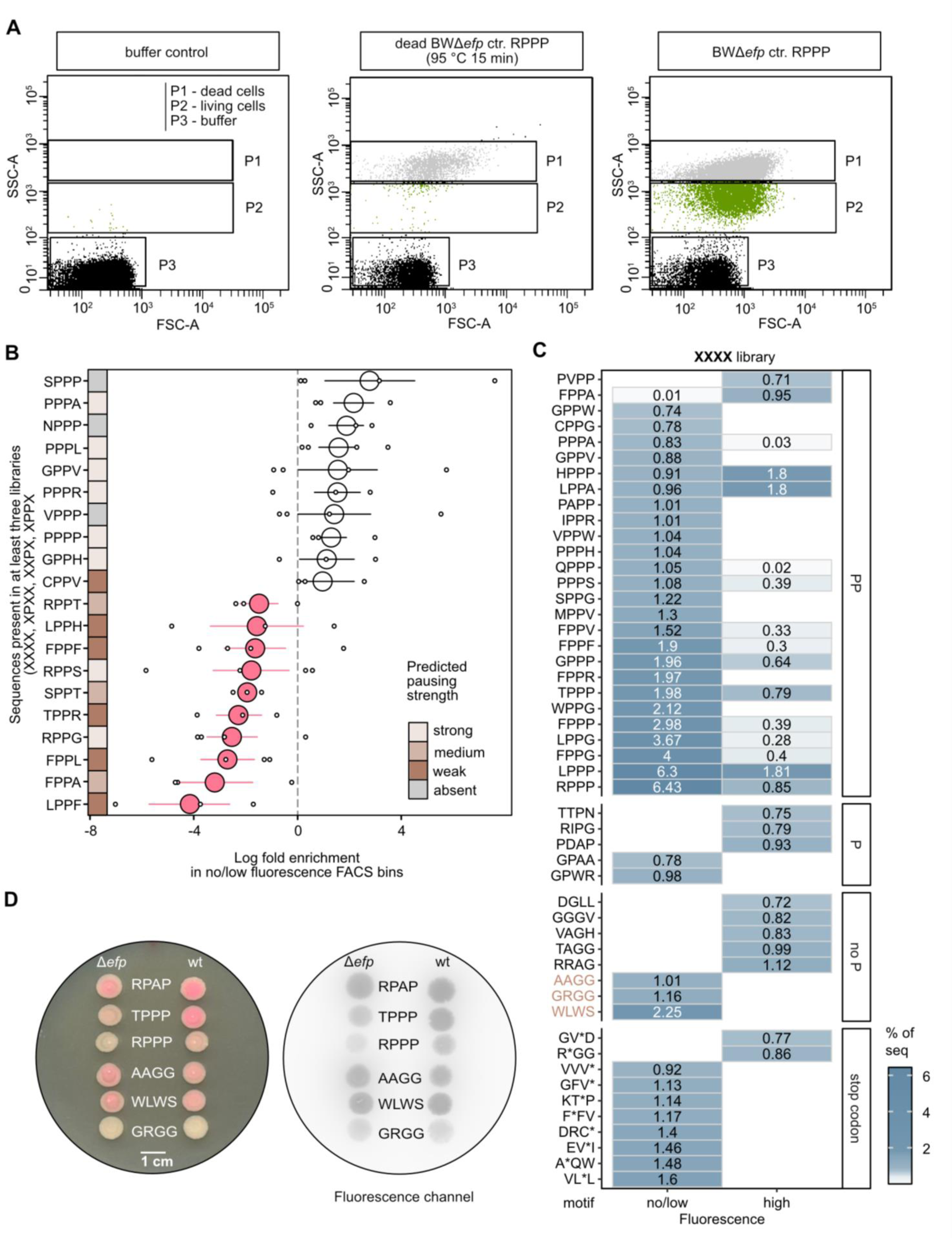
High throughput screening of four motif libraries (XPXX, XXPX, XPPX, XXXX). **A** Gating strategy of living cells by flow cytometry. Living cells (P2) were sorted from the recorded buffer events (P3) and dead cells (P1) considering the Forward Scatter Area (FCS-A) and Side Scatter Area (SSC-A). **B** Summary of the motifs found in at least three out of four libraries (X-X-X-X; X-Pro-X-X; X-X-Pro-X; X-Pro-Pro-X) with the top ten highest and top ten lowest log fold enrichment. Libraries were expressed in *E. coli* BW25113 Δ*efp*. The log fold enrichment was calculated as the natural logarithm of the sequence count found in the no/low fluorescence FACS bin divided by the sequence count found in the high fluorescence FACS bin. Small circles-values in each library, big circles – mean values, lines – standard error of the mean (SEM). Big circles were coloured according to the calculated log fold enrichment value: <0, in pink; >0, in white. Pausing strength predictions were taken from Qi *et al*., 2018^21^. **C** The fifty most frequent amino acid motifs found in the XXXX motif library with the largest count differences between no/low fluorescence and high fluorescence FACS bins. The motifs were divided into categories of motifs containing at least two prolines (PP), one proline (P), no proline (no P) and a stop codon (stop codon, *). X represents all possible amino acids. **D** Translational efficiency measurements in Δ*efp* and wild-type (wt) cells carrying dual reporter plasmids with various motifs, analysed using a spot assay. Motifs analysed: RPAP (Arg-Pro-Ala-Pro), TPPP (Thr-Pro-Pro-Pro), RPPP (Arg-Pro-Pro-Pro), AAGG (Ala-Ala-Gly-Gly), WLWS (Trp-Leu-Trp-Ser) and GRGG (Gly-Arg-Gly-Gly). Right panel shows the fluorescence analysis of the spot assay plate.

A number of studies have reported on ribosomal pausing, which occurs at motifs containing consecutive prolines that can be rescued by EF-P^14–16,21,41^. So far only few studies report about motifs containing no prolines, that depend on EF-P for efficient translation^37,42^. Consequently, we wanted to see whether we could detect any non-proline motifs within the XXXX library that reduced translational efficiency of the fluorophore. We extracted the top fifty sequences with the largest difference in sequence counts between FACS bins and found four groups of motifs: those containing at least two prolines (PP), those containing one proline (P), those containing no proline (no P) and those containing a stop codon (**Figure 3C**). In accordance with our expectations, motifs containing a stop codon were highly abundant in the no/low fluorescence FACS bin (**Figure 3C**). In our system, stop codons cause the rapid termination of fluorophore translation, leading to the production of a non-functional protein lacking fluorescence. We found that certain motifs within the PP section appeared in both FACS bins but with differing abundance, which may be due to the heterogeneity phenomenon (**Supplementary Figure 3D**) and/or codon effects (**Supplementary Figure 4**). However, the majority of motifs in the PP section appeared in the no/low fluorescence FACS bins, consistent with previous studies indicating that the presence of at least two prolines hinders translation^16,21^.

Interestingly, our experimental system identified two motifs containing a single proline (GPAA, GPWR) which decreased the efficiency of fluorophore translation. Additionally, three motifs without a proline (AAGG, GRGG, WLWS) were identified (**Figure 3C**). To verify these results, we transformed the Δ*efp* and wild-type strain with newly generated dual reporter plasmids containing the motifs AAGG, GRGG or WLWS, and spotted cells on agar assay plates. We found that these motifs negatively impact translational efficiency in the Δ*efp* mutant (**Figure 3D**), which is consistent with the observations shown in **Figure 3C**. The spot assay confirmed that the AAGG and WLWS motifs cause an intermediate reduction in translational efficiency of the fluorophore, while the GRGG motif causes a strong reduction in translation efficiency in the Δ*efp* mutant (**Figure 3D**). When EF-P and its modification machinery were present (wt), translation of the fluorophore could be rescued for the control motifs TPPP and RPPP (**Figure 3D**). However, no rescue of fluorophore translation could be detected for the motifs AAGG, GRGG and WLWS (**Figure 3D**), suggesting that additional factors influence translation efficiency at these motifs.

Overall, we find that translational regulation appears to be more complex than previously thought. Consequently, our system offers a suitable platform to get more insights into this phenomenon in any laboratory.

## Discussion

Translational coupling is a naturally occurring event in which the movement of ribosomes translating upstream genes also directs the translation of downstream genes. This mechanism ensures the coordinated and sequential synthesis of proteins involved in shared biochemical pathways or protein complexes, thereby ensuring efficient cellular function and effective resource allocation^50–54^. Translational coupling has been used in experimental studies to identify optimal translation initiation regions (TIRs) and enhance the synthesis of distinct proteins^43,44,55^. In the present study, we used translational coupling in the design of a dual reporter to investigate translational regulation. This system can be effectively adapted for use on agar assay plates, providing a dual output by selecting for fluorescence and viability, allowing uncomplicated screening.

As a component of the dual reporter system, mScarlet-I offer several advantages in plate-based translational efficiency screenings. Translational coupling of mScarlet-I to the resistance gene cassette (*cat*) resulted in the attenuation of *cat,* which improved reporter sensitivity (**Figure 1**). The presence of *cat* alone allowed only coarse distinctions in translational efficiencies — only motifs that are known to cause strong ribosome pausing and motifs that cause no pausing at all could be distinguished (**Supplementary Figure 1A-C**). This suggests that when present alone in the reporter, *cat* ‘neutralises’ chloramphenicol so effectively that small differences in ribosome pausing become indistinguishable. In contrast, the incorporation of mScarlet-I into the reporter allowed finer discrimination between pausing strengths caused by different motifs (**Figure 1**). In addition, the bright fluorescence, good colour contrast with the plate background (**Figure 1G** and **Figure 2B**) and ability to emit light in the visible red spectrum (FPbase ID: 6VVTK)^56,57^ make mScarlet-I a perfect component of the reporter so that measurements could be performed without additional equipment.

Previous studies have demonstrated that in addition to consecutive prolines, sequences upstream and downstream of prolines can also influence translational efficiency in *E. coli*^16,38^. We were able to confirm and reproduce these observations, particularly in the context of downstream sequences (FPPX) in Δ*efp* (**Figure 2**). Motifs with no or intermediate influence on translational efficiency (FPPF, FPPI, FPPV and FPPY) and high influence on translational efficiency (FPPP) (**Figure 2C**) were consistent with previously published proteome and ribosome profiling data^16,39^. However, we do not exclude the possibility that sequences further away from the motif, as well as the codon context of the motif, might also influence translational efficiency. This should be investigated in future studies.

Our investigations of motifs within the XXXX libraries, conducted using flow cytometry and high-throughput screening, identified new motifs containing one (GPAA, GPWR) or no prolines (AAGG, GRGG, WLWS) that affect translational efficiency (**Figure 3C and D**). Two of the motifs we identified were found to contain more than one glycine. Previous studies have demonstrated that sequences rich in glycine have the potential to cause ribosome pausing^37,58^. mRNA sequences that are similar to the Shine-Dalgarno (SD) sequence can trigger an SD-aSD (Shine-Dalgarno - anti-Shine-Dalgarno) interaction, which in turn causes ribosome pausing^5,59,60^. Nevertheless, it is still unclear how glycine codons affect translational pausing^61^, as some studies suggest that internal SD sequences have no/low effect on translational speed^62,63^. The present study provides a platform that can support future research aimed at clarifying the influence of glycine on translational pausing in different sequence contexts.

In addition to EF-P, the ABCF ATPase YfmR has recently been shown to be involved in the translational rescue of polyproline motifs^64,65^. The reporter described here could easily be adapted to study the full spectrum of motifs rescued by this system in any laboratory. Additionally, while the spectrum of motifs that cause EF-P dependent ribosome pausing has been studied extensively, these experiments have primarily focused on *E. coli* and its β-lysylated post-translationally modified EF-P^16,21,39^. Our recent study showed differences in motif rescue preferences between wild-type *E. coli* and a mutant strain in which the native EF-P was replaced with an unmodified PGKGP subfamily EF-P^36^. This suggests that there is a subset of motifs that are preferentially rescued by different EF-P types. If so, different EF-P types could drive the evolution of bacterial genomes towards motifs that they preferentially rescue, particularly in highly expressed proteins where maximizing the translation rate is essential^21,66^. Such motif bias spectra could be studied with our reporter system.

## Material and Methods

### Bacterial growth

All bacterial strains used in this study were cultivated in lysogenic broth (LB) under agitation at 37°C. If required, growth media was solidified by the addition of 1.5 % (w/v) agar. The following antibiotics were used in this study: carbenicillin sodium salt (100 µg/ml), chloramphenicol (default concentration 34 μg/mL; reporter assays: 1.7 μg/mL, 2.3 μg/mL, 3.4 μg/mL and 3.8 μg/mL).

### Molecular cloning

All strains, plasmids and DNA oligonucleotides used are listed in Supplementary Data 1. Kits, enzymes and polymerases were used according to the manufacturer’s instructions. Q5^®^ High-Fidelity DNA Polymerase (New England BioLab) and One*Taq*^®^ DNA Polymerase were used for PCR amplification. DNA oligonucleotides were ordered from Merck. Amplified genes coding for the fluorophores (*sfgfp* and *mscarlet-I*) and the antibiotic resistance cassette (*chloramphenicol acetyltransferase*, CAT) were linked together with a hairpin loop (weak coupling)^43^ by overlap PCR. DNA was isolated and purified with High-Yield PCR Cleanup and Gel Extraction Kit (Sued Laborbedarf Gauting). All DNA restriction digests were carried out using restriction enzymes in rCutSmart^®^ buffer (New England BioLab). All ligations were performed with the T4 DNA ligase (New England BioLab) at 25 °C for 2 hours. Plasmids were isolated with the Hi Yield Plasmid Mini Kit (Sued Laborbedarf Gauting) and stored at −20°C. Plasmids were verified by colony PCR and sanger sequencing. All nucleotide sequences were analyzed with CLC Main Workbench version 8.1.2 (Qiagen).

### Growth and fluorescence measurements in 96-well plates

To find the optimal concentration of chloramphenicol for the pausing strength measurements with the dual reporter, serial dilutions were tested. Plasmids pBAD24_motif_*sfgfp*_*cat* and pBAD24_motif_*mscarlet-I*_*cat* (motif: RPAP, TPPP or RPPP) were independently transformed into *E. coli* BW25113 Δ*efp* and incubated overnight at 37 °C on LB agar plates supplemented with 100 µg/ml carbenicillin sodium salt. Single colonies were inoculated into fresh LB supplemented with 100 µg/ml carbenicillin sodium salt and incubated overnight at 37°C under constant shaking (180 rpm). The overnight cultures were reinoculated into fresh LB supplemented with 100 µg/ml carbenicillin sodium salt and grown until reaching an absorption of 0.3 at 600 nm (OD_600_). Each culture was transferred to a final OD_600_ of 0.01 into a 96-well plate, containing 0.2 % (w/v) arabinose and different concentrations of chloramphenicol (1.7 μg/mL, 2.3 μg/mL, 3.4 μg/mL, 3.8 μg/mL) with a final volume of 200 μL/well. Plates were incubated overnight at 37 °C under constant shaking (150 rpm). Endpoint OD_600_ was determined with Tecan Infinite 200 Pro (number of flashes: 25). Fluorescence was measured with GE Typhoon Trio Imager (laser: green, 532 nm; emission filter: 580 BP 30 Cy3; photo-multiplier tube, PMT: 450).

For all time-course fluorescence and growth measurements, the cells were grown for 20 hours in M9 minimal media^67^ (composition: 33.7 mM Na_2_HPO_4_, 22 mM KH_2_PO_4_, 8.55 mM NaCl, 9.35 mM NH_4_Cl, 1 mM MgSO_4_, 0.3 mM CaCl_2_, 1 μg biotin, 1 μg thiamin, trace elements, 0.4 % [w/v] glucose) prior to the start of the measurements in the plate reader. Growth and fluorescence were monitored in 10 min intervals for 35 hours at 37 °C with Tecan Infinite 200 Pro (excitation wavelength: 565 nm; emission wavelength: 594 nm; gain: 50; number of flashes: 25; shaking between measurements: 180 rpm, orbital).

EF-P complementation assay plates were incubated for 24 to 34 hours until fluorescence measurements.

### Generation of XPPX, PXXX, XPXX, XXPX, XXXP, XXXX libraries

The forward primers were designed to contain six to twelve degenerated nucleotides that code for all possible codons. The fixed prolines in the motif were coded by CCG codon. Degenerated codons were designed to appear fifteen nucleotides downstream of the start codon of the fluorophore gene. Libraries were amplified using the degenerated forward primer and reverse primer with complementary regions to the forward primers to enable plasmid circularization afterwards (**Figure 3A**). The PCR program used for library amplification was as follows: initial denaturation, 98 °C for 30 seconds (sec); amplification (35 cycles), 98 °C for 10 sec followed by 57°C for 30 sec and 72 °C for 180 sec; final extension, 72 °C for 120 sec. 40 µL of the PCR product was treated with *Dpn*I. Chemically competent *E. coli* MC1061 was transformed with the digested and purified PCR products to enable plasmid circularization by homologous recombination^43,44,47^. The transformation was transferred to a flask with 15 mL LB, supplemented with 100 µg/ml carbenicillin sodium salt, and incubated overnight at 37 °C under constant shaking (180 rpm). The libraries were isolated with the Hi Yield Plasmid Mini Kit (Sued Laborbedarf Gauting).

### Screening of libraries on assay plates

The final libraries were prepared according to the protocol described by Rennig *e*t al.^43^ and Shilling *et* al.^44^, with certain modifications. Chemically competent *E. coli* BW25113 Δ*efp* was transformed with 500 ng of the library with a 1-hour recovery phase in fresh LB without antibiotics. The transformants were inoculated into 3 mL fresh LB containing 100 µg/ml carbenicillin sodium salt and incubated overnight at 37 °C under constant shaking (180 rpm). 30 µL of this overnight culture was reinoculated into 3 mL fresh LB (containing 100 µg/ml carbenicillin sodium salt) and incubated for another round overnight at 37 °C under constant shaking (180 rpm). 100 µL of this overnight culture was inoculated into fresh 5 mL LB supplemented with 34 μg/mL chloramphenicol and grown until reaching an OD_600_ of 0.3. 10^3^ - 10^5^ cells were plated on assay agar plates, supplemented with 3.4 μg/mL chloramphenicol and 0.2 % (w/v) arabinose. Plates were incubated for 16 hours at 30 °C and kept afterwards for ∼ 120 hours at 25°C. Fluorescence was measured with GE Typhoon Trio Imager (laser: green, 532 nm; emission filter: 580 BP 30 Cy3; pmt: 450).

### Microscopy

To analyze the effect of different motifs on the production of mScarlet-I, chemically competent *E. coli* BW25113 Δ*efp* was transformed with the plasmids pBAD24_motif_*mscarlet-I_cat* (motif: RPAP, TPPP or RPPP). Cells carrying the plasmids were cultivated overnight at 37 °C under constant shaking (180 rpm) in LB supplemented with 100 µg/ml carbenicillin sodium salt. The overnight cultures were inoculated into fresh LB supplemented with 100 µg/ml carbenicillin sodium salt and grown at 37 °C under constant shaking (180 rpm) until reaching an OD_600_ of 0.15. 0.2 % (w/v) arabinose was then added to the cultures and incubation continued for 24 hours. All cells were washed in phosphate buffered saline (PBS; 93.6 mM NaCl, 2.7 mM KCl, 10 mM Na_2_HPO_4_ * 2 H_2_O, 2 mM KH_2_PO_4_) and 2 μL of a culture with an OD_600_ of 0.5 were spotted on a 1 % (w/v) agarose pad (in PBS) and covered with a coverslip. Microscopic pictures were taken using the Leica DMi8 inverted microscope equipped with a Leica DFC365 FX camera (Wetzlar Germany). An excitation wavelength of 546 nm and a 605 nm emission filter with a 75-nm bandwidth were used for mScarlet-I fluorescence with an exposure of 20 ms, gain 1, and 100% intensity. To quantify the relative fluorescent intensities (RF) of single cells, phase contrast and fluorescent images were analyzed using the ImageJ^68^ plugin MicrobeJ^69^. Default settings of MicrobeJ were used for cell segmentation (Fit shape, rod-shaped bacteria) apart from the following settings: area: 0.1-max µm^2^; length: 1.2-5 µm; width: 0.1-1 µm; curvature 0.-0.15 and angularity 0.-0.25 for *E. coli* cells. In total 516 cells were quantified per strain. Background of the agarose pad was subtracted from each cell per field of view.

### Flow cytometry and FACS

The influence of different motifs on the translational efficiency of the fluorophore mScarlet-I was analyzed using high throughput screening with flow cytometry and fluorescence associated cell sorting (FACS). Cells containing the dual reporter plasmid were grown in LB supplemented with 100 µg/ml carbenicillin sodium salt and 0.2 % (w/v) arabinose for 16 hours at 37 °C under constant shaking (180 rpm). The overnight incubation was necessary to provide enough time for mScarlet-I to mature before the FACS analysis. Cells were washed in PBS and seeded to an OD of 0.1 (10^8^ cells) in 10 mL PBS (4°C) prior to the measurements. For the dead cells control, the culture was boiled at 95 °C for 15 min. The gating strategy for living cells is depicted in **Figure 4A**. Flow cytometry measurements and FACS were performed using FACSAria^TM^FusionII Cell Sorter (BD Biosciences) with the following settings: 438 V(FSC), 277 V (SSC), 490 V (PE-Texas Red). Red fluorescence intensities were quantified using PE-Texas Red and displayed on a standard logarithmic scale. Flow cytometry data was processed with FACSDiva Software (BD Biosciences). Motifs found in cells with no/low fluorescence (intensity below 10^2.1^ on the logarithmic scale) were classified as those capable of slowing down the translational efficiency of the fluorophore. In contrast, motifs found in cells with high fluorescence (intensity above 10^3.1^ on the logarithmic scale) were classified as those having only a marginal influence on the translational efficiency of the fluorophore.

### Library preparation for Illumina high throughput sequencing

Plasmid from each FACS bin were isolated with the Hi Yield Plasmid Mini Kit (Sued Laborbedarf) and served as template DNA for Illumina-specific amplification required for sequencing^70^. Custom made DNA oligos were designed (Merk), consisting of 21 bp Illumina primer sequence, distinct barcoding tag^70^ and a sequence complementary to the dual reporter plasmid (forward primer: 81 bp upstream of the library motif, reverse primer: 167 bp downstream of the library motif; **Supplementary Figure S5**). Purified PCR products were loaded on a 1% (w/v) agarose gel to estimate the amount of each amplicon and pooled in equimolar concentrations^70^. The library was finalized for sequencing according to the manufacturer’s protocols (http://supportres.illumina.com/documents/documentation/system_documentation/miseq/ preparing-libraries-for-sequencing-on-miseq-15039740-d.pdf). Paired-end sequencing of 2 x 300 bp (base pairs) with two additional 8 bp index reads was performed on an Illumina MiSeq platform (Illumina Inc.) with v3 chemistry. All Sequencing data generated in this study is deposited in the European Nucleotide Archive (ENA) at EMBL-EBI under accession number PRJEB77738

### Bioinformatic analysis of Illumina library sequencing

Sequences were first quality filtered using FASTQ version 0.23.4^71^. Sequences were trimmed using a dynamic approach—if the mean quality of bases within a sliding window of 4 nucleotides dropped below a cutoff of 20, sequences were trimmed starting at the first base within the window (`--cut_right`). Paired reads were corrected using the overlapping region (`--correction`). Next, PANDAseq version 2.11 was used with default parameters to merge paired end sequences^72^. SABRE version 1.0 was used to demultiplex sequences^73^, and custom python scripts were used to extract the sequence region containing the library. The R package Vegan version 2.6 was used to rarefy sequences to the level of the sample with the lowest sequence count (this level varied depending on the samples being compared but was never lower than 98335 sequences)^74^. All statistical analyses on the sequence data were done in R version 4.3.2^75^, and all related figures were created with ggplot2 version 3.4.4^76^.

### Statistical analysis and reproducibility

All measurements are from at least three biological replicates except the plate-based library screening (**Figure 2C**, the count of found motifs is indicated in the figure) and FACS. Data comparisons were done using the student’s unpaired two-sided t test (Microsoft Office Excel 2021) and are described in each figure legend. Error bars in bar graphs represent the standard deviation (SD). Values with a calculated p *value* below 0.05 were considered as significantly different.

## Acknowledgements

We thank Dr. Andreas Brachmann and the Genomics Service Unit at the LMU for fruitful discussions and excellent support in the sequencing of the Illumina libraries. We thank the Core Facility Flow Cytometry at Biomedical Center Munich for the great technical support. We thank Lukas Voshagen for great technical assistance. This work was financially supported by the Deutsche Forschungsgemeinschaft (DFG, German Research Foundation) grant JU270/20-1, project number 449926427, and RTG2062 (Molecular Principles of Synthetic Biology) to K.J.

## Author contributions

U.T. and K.J. designed the research; U.T. and K.B. performed the research; U.T., T.B. and S.B. analysed the data; U.T. and K.J. wrote the manuscript; K.J. funding acquisition.

