## Supplementary Information for "A versatile dual reporter to identify ribosome pausing motifs alleviated by translation elongation factor P"

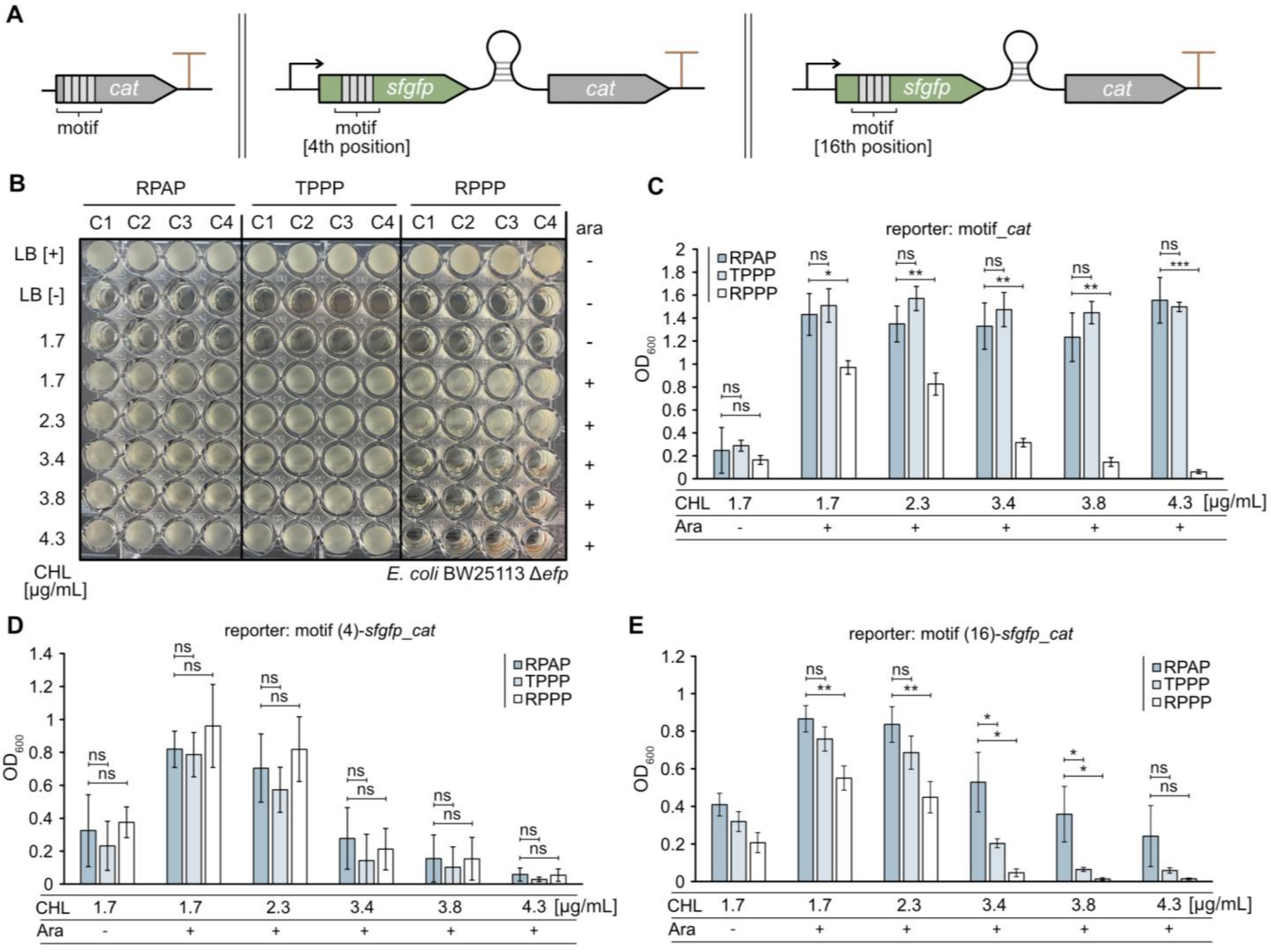


**Supplementary Figure 1. Bacterial survival-based determination of translational efficiencies by using a heterologous dual reporter system. A** Construction of reporters for translational efficiency measurements. The reporters were designed to consist of either one reporter gene encoding an antibiotic resistance protein (left panel; chloramphenicol acetyltransferase, CAT) or two reporter genes encoding a fluorophore sfGFP and CAT, linked with a weak translational coupling device[1]. The motifs were located either from the fourth (4th) or from the sixteenth (16th) nucleotide position of the gene coding for the fluorescent protein. The arrows illustrate promoter locations. **B, C** 96-well assay plate of the growth measurements (**B**) and growth quantification (**C**) of *E. coli* BW25113 Δ*efp*. The strain contains the reporter plasmid pBAD24_motif_*cat*. **D, E** Growth quantification of *E. coli* BW25113 Δ*efp* containing the dual reporter plasmid pBAD24_motif_*sfgfp_cat* with the motif at the 4th (**D**) or the 16th (**E**) nucleotide position of the gene sequence coding for fluorescent protein. RPAP, TPPP and RPPP motifs were used to assess no stalling/ stalling of the ribosome (Figure 1A). Error bars indicate the standard deviation (SD) of three independent biological replicates. Statistics: student's unpaired two-sided t test (****$p$ < 0,0001; ***$p$ < 0.001; **$p$ < 0.01; *$p$ < 0.05; ns $p$ > 0.05). 1.7 µg/mL CHL and no Ara (RPAP vs TPPP: ns $p$ = 0.750 [C], ns $p$ = 0.570 [D]; RPAP vs RPPP: ns $p$ = 0.524 [C], ns $p$ = 0.732 [D]); 1.7 µg/mL CHL and 0.2 % (w/v) Ara (RPAP vs TPPP: ns $p$ = 0.589 [C], ns $p$ = 0.752 [D], ns $p$ = 0.099 [E]; RPAP vs RPPP: *$p$ = 0.017 [D], ns $p$ = 0.423 [D], **$p$ = 0.001 [E]); 2.3 µg/mL CHL and 0.2 % (w/v) Ara (RPAP vs TPPP: ns $p$ = 0.092 [C], ns $p$ = 0.399 [D], ns $p$ = 0.093 [E]; RPAP vs RPPP: **$p$ = 0.004 [C], ns $p$ = 0.513 [D], **$p$ = 0.002 [E]); 3.4 µg/mL CHL and 0.2 % (w/v) Ara (RPAP vs TPPP: ns $p$ = 0.359 [C], ns $p$ = 0.381 [D], *$p$ = 0.036 [E]; RPAP vs RPPP: **$p$ = 0.003 [C], ns $p$ = 0.634 [D], *$p$ = 0.013 [E]), 3.8 µg/mL CHL and 0.2 % (w/v) Ara (RPAP vs TPPP: ns $p$ = 0.184 [C], ns $p$ = 0.646 [D], *$p$ = 0.041 [E]; RPAP vs RPPP: **$p$ = 0.002 [C], ns $p$ = 0.991 [D], *$p$ = 0.027 [E]); 4.3 µg/mL CHL and 0.2 % (w/v) Ara (RPAP vs TPPP: ns $p$ = 0.649 [C], ns $p$ = 0.311 [D], ns $p$ = 0.147 [E]; RPAP vs RPPP: ***$p$ = 0.001 [C], ns $p$ = 0.925 [D], ns $p$ = 0.095 [E]). Ara – arabinose, C1-C4 – clone number, CHL – chloramphenicol, *msc - mscarlet-I, sfgfp - super folded green fluorescent protein.*

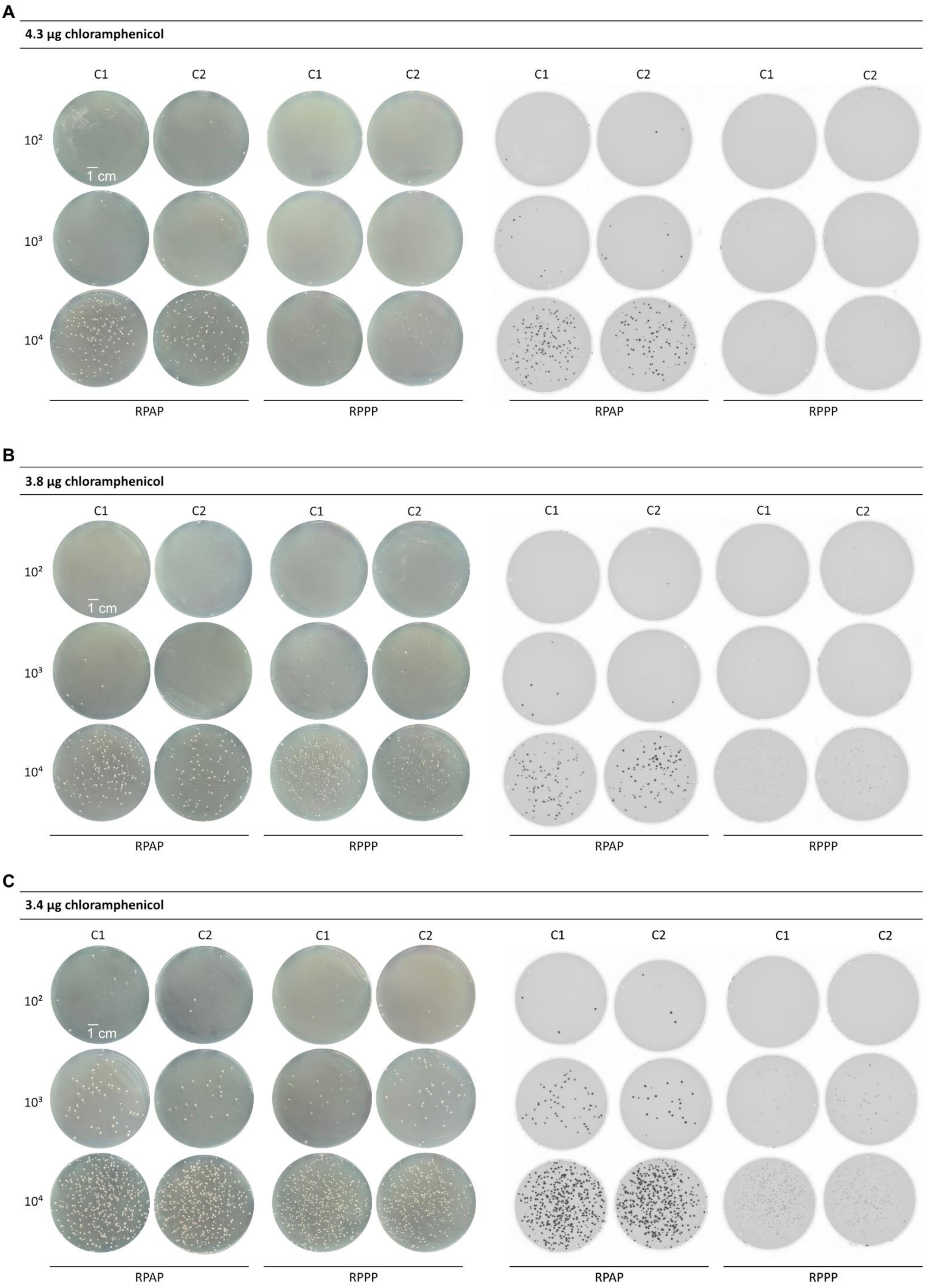

A
4.3 µg chloramphenicol
C1
C2
C1
C2
C1
C2
C1
C2
10²
10³
10⁴
1 cm
RPAP
RPPP
RPAP
RPPP
B
3.8 µg chloramphenicol
C1
C2
C1
C2
C1
C2
C1
C2
10²
10³
10⁴
1 cm
RPAP
RPPP
RPAP
RPPP
C
3.4 µg chloramphenicol
C1
C2
C1
C2
C1
C2
C1
C2
10²
10³
10⁴
1 cm
RPAP
RPPP
RPAP
RPPP

**Supplementary Figure 2. Influence of the chloramphenicol concentration and the motif on the survival and fluorescence of the reporter strain. A-C** Growth (left panel) and colony fluorescence (right panel) analysis of *E. coli* BW25113 Δ*efp* on LB agar assay plates, supplemented with 0.2 % (w/v) arabinose and 4.3 µg (**A**), 3.8 µg (**B**) or 3.4 µg (**C**) chloramphenicol. $10^2$ – $10^4$ of cells were plated. The strain was transformed with the dual reporter plasmid pBAD24_motif_*mscarlet-I_cat*, containing either RPAP (Arg-Pro-Ala-Pro) or RPPP (Arg-Pro-Pro-Pro) amino acid motif. C1 – C2 – clone number.

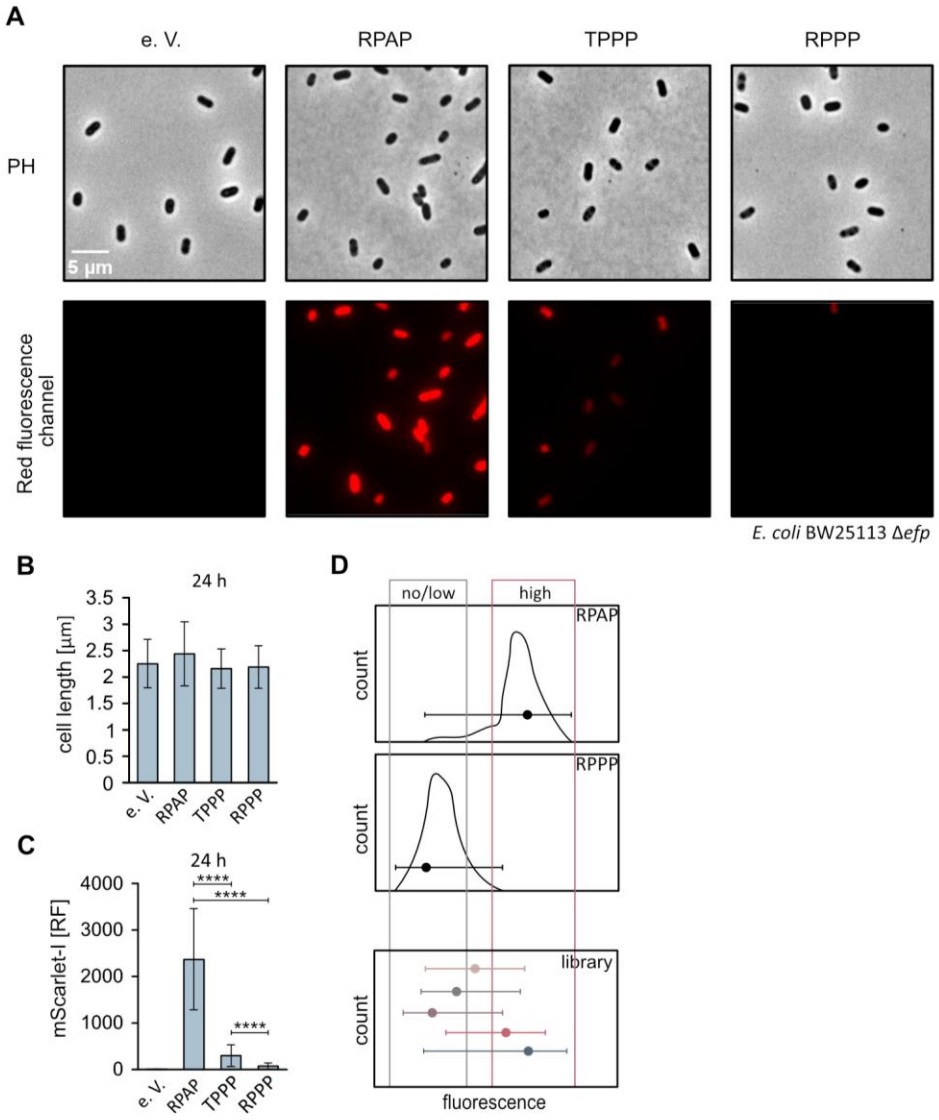


**Supplementary Figure 3. Heterogeneity in plasmid-based fluorophore expression. A** Fluorescence microscopic images of BW25113 Δ*efp* transformed with dual reporter plasmid pBAD24_motif_*mscarlet-I_cat*. The top panel shows phase contrast (PH) images, the bottom panel - images from the red fluorescent channel. RPAP (Arg-Pro-Ala-Pro), TPPP (Thr-Pro-Pro-Pro) and RPPP (Arg-Pro-Pro-Pro) amino acid motifs were used to asses no stalling/ stalling of the ribosome. **B** Bar graph showing the average bacterial cell length from a total of 516 cells containing the reporter with the indicated amino acid motif. **C** Quantification of relative cell fluorescence from microscopic images. Microscopic images were taken 24 h after induction with 0.2 % (w/v) arabinose. **D** Schematic representation of the heterogeneity phenomenon in flow cytometry. e. V. – empty vector, h – hour, RF – relative fluorescence. Error bars indicate the standard deviation (SD) with following statistics: student's unpaired two-sided t test (****$p$ < 0,0001; ***$p$ < 0.001; **$p$ < 0.01; *$p$ < 0.05; ns $p$ > 0.05). RPAP vs TPPP: ****$p$ < 0,0001; RPAP vs RPPP: ****$p$ < 0,0001; TPPP vs RPPP: ****$p$ < 0,0001 [**C**].

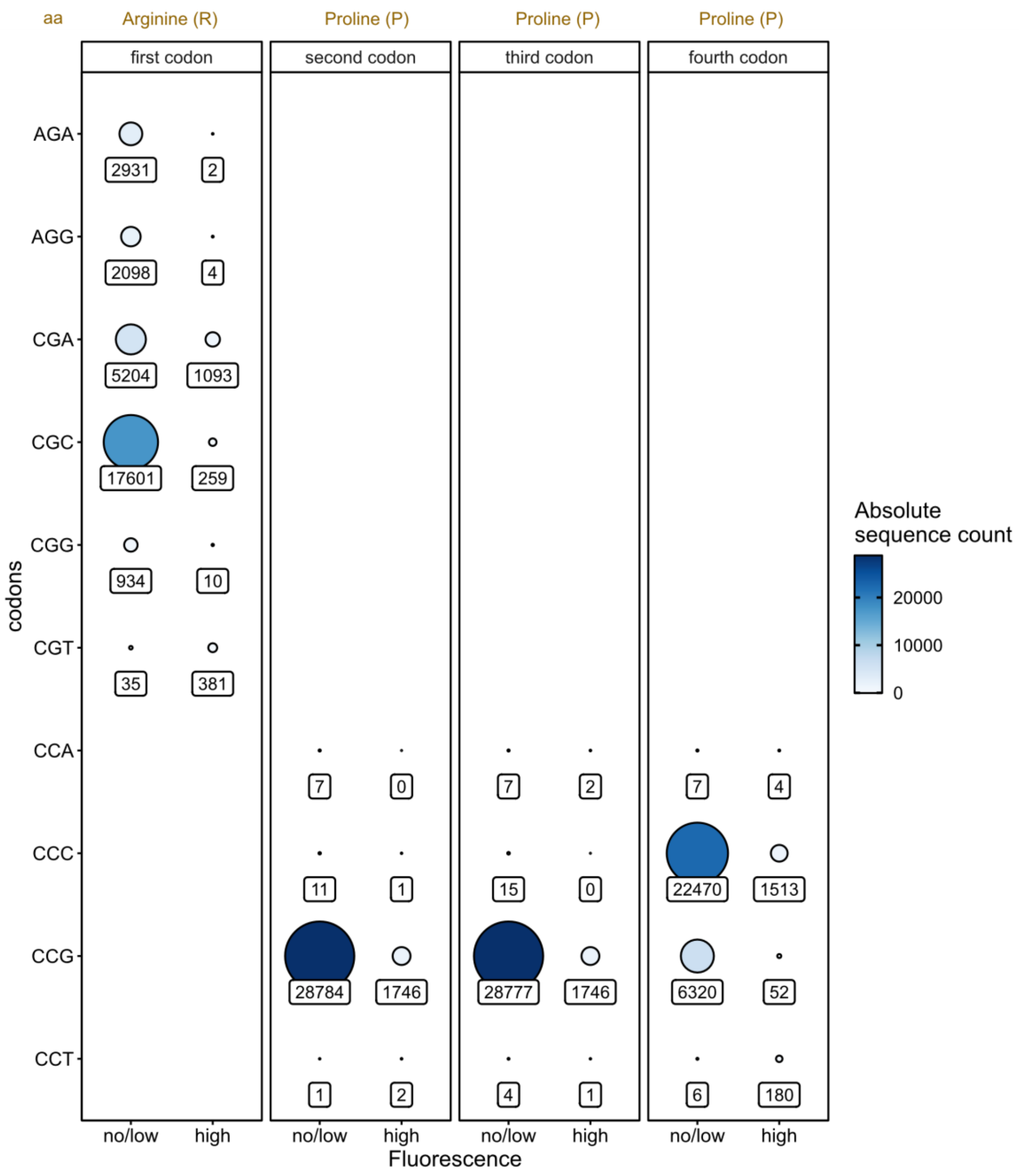


**Supplementary Figure 4. Distribution of codons encoding the motif RPPP in the library XXXX.** Absolute sequence counts are depicted in numbers. The size of the circles corresponds to the count magnitude. aa – amino acid.

| | Illumina primer sequence | barcoding tag | complementary sequence to target DNA |
|---|---|---|---|
| ilumina_seq_F1 | TACACGACGCTCTTCCGATCT | TCAT | TTAGCGGATCCTACCTGACG |
| ilumina_seq_F2 | TACACGACGCTCTTCCGATCT | AAGTGA | TTAGCGGATCCTACCTGACG |
| ilumina_seq_F3 | TACACGACGCTCTTCCGATCT | TGCGAGA | TTAGCGGATCCTACCTGACG |
| ilumina_seq_F4 | TACACGACGCTCTTCCGATCT | GACATCCA | TTAGCGGATCCTACCTGACG |
| ilumina_seq_R2 | CAGACGTGTGCTCTTCCGATCT | CGCTCA | ACATGAACTGAGGGGACAGG |
| ilumina_seq_R3 | CAGACGTGTGCTCTTCCGATCT | GCTAACA | ACATGAACTGAGGGGACAGG |
| ilumina_seq_R4 | CAGACGTGTGCTCTTCCGATCT | TTGACCAG | ACATGAACTGAGGGGACAGG |

**Supplementary Figure 5. Primer design for Illumina library sequencing.** Forward primers (F1-F4) and reverse primers (R2-R4) were designed to contain a sequence required for Illumina sequencing, a barcoding tag[2] for sample distinction, and a sequence complementary to the target DNA (library plasmids).
